# High-throughput quantitative screening of glucose-stimulated insulin secretion and insulin content using automated MALDI-TOF mass spectrometry

**DOI:** 10.1101/2023.02.14.528514

**Authors:** Clément Philippe Delannoy, Egon Heuson, Adrien Herledan, Frederik Oger, Bryan Thiroux, Mickaël Chevalier, Xavier Gromada, Laure Rolland, Philippe Froguel, Benoit Deprez, Sebastien Paul, Jean-Sébastien Annicotte

**Affiliations:** Univ. Lille, Inserm, CHU Lille, Institut Pasteur de Lille, CNRS, U1283 – UMR 8199 – EGID, F-59000, Lille, France; Univ. Lille, CNRS, Centrale Lille, Univ. Artois, UMR 8181 – UCCS- Unité de Catalyse et de Chimie du Solide, F-59000, Lille, France; Univ. Lille, Inserm, Institut Pasteur de Lille, U1177 – Drugs and Molecule for Living Systems, F-59000, Lille, France; Univ. Lille, UMRt BioEcoAgro 1158-INRAE, équipe Métabolites secondaires d’origine microbienne, Institut Charles Viollette, F-59000 Lille, France; Univ. Lille, Inserm, CHU Lille, Institut Pasteur de Lille, U1167 - RID-AGE - Facteurs de risque et déterminants moléculaires des maladies liées au vieillissement, F-59000 Lille, France

## Abstract

Type 2 diabetes (T2D) is a metabolic disorder characterized by loss of pancreatic β- cell function, decreased insulin secretion and increased insulin resistance, that affects more than 400 million people worldwide. Although several treatments are proposed to patients suffering from T2D, long-term control of glycemia remains a challenge. Therefore, identifying new potential drugs and targets that positively affect β-cell function and insulin secretion remains crucial. Here, we developed an automated approach to allow the identification of new compounds or genes potentially involved in β-cell function in a 384-well plate format, using the murine β-cell model Min6. Using MALDI-TOF mass spectrometry, we have implemented a high-throughput screening (HTS) strategy based on the automation of a cellular assay allowing to detect insulin secretion in response to glucose, quantitative detection of insulin, in a miniaturized system. As a proof of concept, we screened siRNA targeting well-know β-cell genes and 1600 chemical compounds and identified several molecules as potential regulators of insulin secretion and/or synthesis, demonstrating that our approach allows HTS of insulin secretion *in vitro*.

## Introduction

Type 2 Diabetes (T2D) is characterized by high blood glucose levels and develops due to inadequate pancreatic β-cell function (i.e. insulin secretion) and peripheral insulin resistance. In Europe, the global prevalence of diabetes is currently estimated at 8 % of the population, with T2D representing 90 % of cases [1]. Although lifestyle modification is the first reference treatment for T2D, modifying the patient’s habits is often ineffective to stabilize glycemia and combinations of pharmacological treatments (metformin, sulfonylurea, incretin enhancers, GLP-1 mimetics, etc…) are often re-quired to treat T2D [2,3]. Yet, long-term control of glycemia remains a challenge for most patients, particularly when β-cell capacity to secrete insulin decreases with age [4].

In the context of research aimed at discovering new therapeutic targets for T2D, the identification of new natural or synthetic compounds or target genes that have the capacity to restore or increase insulin secretion is essential. Therefore, developing high-throughput screening (HTS) strategies to rapidly measure insulin secretion or synthesis is crucial to address the unmet need of new efficient molecules for T2D patients. Currently, several strategies have been developed based on assays aiming at measuring secreted hormones or specific proteins (e.g. insulin or c-peptide)[5], mainly through Enzyme-Linked Immuno Sorbent Assay (ELISA) approaches. However, this technique, although widely used in academic research and clinical diagnosis, has several limitations such as the technical detection of the target protein, its precise quantification, the duration of the experiment and the costs of the reagents. Altogether, these limitations preclude the use of ELISA strategies for real HTS approaches to evaluate the action of a large collection of potential drugs that can stimulate insulin secretion.

Along with ELISA, complementary techniques for high-throughput screening of insulin secretion have recently emerged with high potential for T2D therapy. To overcome the experimental and technical limits of ELISA, several laboratories have developed genetically-engineered cellular tools to measure β-cell function. In these modified cells, the use of immunofluorescence microscopy or luciferase-based approaches allow the measurement of a modified C-peptide [5–9]. Although easy to handle, these genetically-modified tools partially address the functionality of the β cell since they do not directly assess the level of production of endogenous insulin content or the secretion of insulin in response to a physiological stimulation such as glucose. Therefore, to tackle these limitations, we developed a new strategy aiming at implementing an automated approach that could allow HTS of drugs that efficiently modulate glucose-stimulated insulin secretion. This automated process, using mass spectrometry to directly measure insulin, is not only faster but also less expensive, enabling for the first time the implementation of high-throughput exploratory strategies to identify new biological targets or bioactive compounds through the screening of chemical or siRNA libraries. Here, we describe a high-throughput screening approach based on Matrix-Assisted Laser Desorption/Ionization-Time of Flight-mass spectrometry (MALDI-TOF) to quantify insulin production and secretion in a mouse model of β cells, the Min6 cell line. We automated, in a 384-well format, siRNA-based loss-of-function of candidate genes to study their effect on insulin secretion. Finally, as a proof of concept, we screened more than 1600 chemical compounds and identified several drugs that modulate insulin secretion or content.

## Materials and Methods

### Cell culture and treatments

MIN6 cells (Addexbio) were maintained in 25 mM Glucose, Glutamax DMEM medium (Gibco, 31966-021), supplemented with 15 % heat-inactivated fetal bovine, 100 μg/ml penicillin-streptomycin and 55 μM beta-mercaptoethanol (Gibco) and cultured in a humidified atmosphere with 5 % CO_2_ at 37 °C. Cells were seeded at 20,000 cells/well using a Multidrop Combi dispenser (Thermo Fisher Scientific, Waltham, MA) into 384-well plate black *µ*clear. Cells were then treated with different chemical compounds and/or siRNA as described below (see Supplementary table 1 for the list of siRNA used in this study).

### Automated Glucose-stimulated insulin secretion (GSIS) in 384-well format

For GSIS experiments, Min6 cells were plated in 384-well plate (2.10^4^ cells/well) and were incubated in 80 µL of starvation buffer (Krebs Ringer buffer (KRB) supplemented with 0.5 % BSA for 1 h at 37 °C and 5 % CO_2_. The automated GSIS pipeline protocol started on the BioCel platform system (Agilent) including integrated devices for incubation, washing, distribution, pipetting liquid handling system, microplate stacking or sealing systems. All the tasks are sequentially operated on each device using the Direct Drive Robot (DDR) arm (Agilent Biotechnologies) positioned at the center of the platform. At the beginning of the run, microplates placed at 37 °C in 5 % CO_2_ atmosphere in the Incubator (Liconic) are conveyed via a telescopic lift arm on the deck of the platform. The cells are first washed from their culture medium by 4 washes of KRB-BSA with the EL406 washer distributor liquid handling device (Biotek) suitable for 384-well plates. A control of the dispense (with tubes angled to 20°) and aspiration of liquid due to the dual-action manifold system of the device permitted to reduce completely the loose of cells layers during this washing step. The cells are then incubated for 1 hour at 37 °C in KRB-BSA for the starvation stage. To avoid BSA interferences in MALDI-TOF mass spectrometry analysis, Min6 cells are washed 5 times with BSA-free KRB buffer supplemented with 2.8 mM glucose after starvation using a Biotek washing robot. After 1 h at 37 °C, cells are washed 5 times with KRB buffer and supplemented with 2.8 mM glucose by distribution of a solution of KRB-Glucose using the disposable peristaltic pump cassette system (EL406 washer distributor), for 1h at 37 °C and 5 % CO_2_. 100 µl of the supernatant were collected for insulin quantification in 2.8 mM glucose condition using a Bravo liquid handler and cells were subsequently incubated in 80 µL of KRB buffer containing 20 mM glucose for 1h at 37 °C and 5 % CO_2_. 100 µl of the supernatant were collected for insulin quantification in 20 mM glucose condition. The intracellular insulin content was recovered in 40 µL of lysis buffer containing 75 % ethanol and 1.5 % HCl. Microplates with all collected samples are sealed with PlateLoc thermal Microplate Sealer (Agilent Technologies) after collecting step with Bravo and frozen at -80°C until analysis. Insulin concentration was measured through ELISA according the manufacturer’s instructions (Mercodia) or mass spectrometry as described below. Samples were frozen at -80 °C before further processing.

### Automated siRNA reverse transfection in 384-well format

To validate a potential functional effect on GSIS, we selected two siRNA targeting genes previously known to control insulin secretion (ON-TARGETplus SMARTpool siRNA targeting mouse Pdx1 (L-040402-01-0005) and mouse Kcnj11 (L-042183-00-0005)) and two controls (non-targeting negative controls and siGLO fluorescent control). siRNA transfection was performed by reverse transfection using 0.375 % Dharmafect1 transfection reagent (GE Dharmacon) and 50 nM siRNA. siRNAs (200 nL of a 20 µM stock; 0.16 pmol/well) were dispensed into 384-well assay plates (Greiner Bio-One; 78109) using an Echo 550 Series Liquid Handler (Labcyte). On the day of assay, plates that contained siRNAs were thawed and equilibrated to room temperature. Dharmafect1 transfection reagent (0.3 µL/well in DMEM media, Gibco) was added to the assay plates using a Multidrop Combi dispenser (Thermo Fisher Scientific, Waltham, MA), with a standard tube dispensing cassette. After 20 min of incubation at room temperature, cells (65 µL of 300,000 cells/mL; 20,000 cells/well) were added to the assay plates with the Multidrop Combi dispenser and standard tube dispensing cassette. Assay plates were incubated at 37 °C, 5 % CO_2_ in a controlled-humidity incubator and cultured for 48 h before GSIS assays. Samples were frozen at -80 °C before further processing.

### Automated incubation with chemical compounds

Min6 cells (80 µL of 250,000 cells/mL; 20,000 cells/well) were added to the assay plates with the Multidrop Combi dispenser (ThermoFisher) and standard tube dispensing cassette. Assay plates were incubated at 37 °C, 5 % CO2 in a con-trolled-humidity incubator during 24 h. Repaglinide (Euromedex - R1426) and fiazoxyle (CliniSciences - BG0437) were used at concentration of 100 nM and 100 µM. A library containing 1600 lead-like molecules was obtained from Asinex company (Mos-cow, Russia). The compounds were dissolved at a concentration of 10 mM in DMSO and one batch of this library was distributed in a 384 well LDV microplate (LP-0200, Beckman coulter). For automated screening assay, 200 nL of 1600 Asinex compounds (5 microplates) or reference products were transferred in intermediate microplates (Greiner Bio-One; 78109) from source 384 wells LDV microplate using nanoacoustic transfer device (Echo 550), then stored at -20 °C until used. 18 h before automated GSIS protocol, using the BioCel platform, intermediate microplates were filled with 60 µL of cell culture medium with EL406 washer distributor (Cassette for distribution). Then, cell microplates previously placed in the incubator were sent to the EL406 washer distributor for aspiration step in order to leave a remaining volume of 40 µL per well. On the deck of the Bravo device, compounds diluted in medium in intermediate microplate were mixed 4 times and a volume of 20 µL were transferred in cell microplate for a treatment at the final concentration of 10 µM. Cells were treated overnight with different compounds and samples were frozen at -80 °C before further processing.

### Automated MALDI-TOF mass spectrometry analysis

MALDI target were prepared in an automated way using a Biomek NX^p^ liquid handler (Beckman Coulter, Fullerton, CA). The detailed robot routine is available in the Supplementary Information. First, the 384-wells microplate (Greiner) containing the previously prepared samples was placed onto the robot deck, along with the other required labwares. Then, for cellular content samples only, 40 µL of MS-grade water was added to the 40 µL of samples to dilute it by a factor of 2. The dilution is done by mixing 3 times 10 µL of the resulting solution. Then 30 µL of a stock solution of bovine insulin (Sigma-Aldrich) at 10 μg.mL^-1^ in 3 mM hydrochloric acid (pH 2.5) (for content samples, 5 μg.mL^-1^ for extracellular high and low glucose samples) was reparteed into a 2^nd^ 384-wells microplate. In parallel, 15 µL of the MALDI matrix solution containing sinapic acid (Sigma-Aldrich) at 10 mg.mL^-1^ in 50/49/1:acetonitrile/MS-grade water/trifluoroacetic acid (Sigma Aldrich) was added in each well of a 3^rd^ separated 384-wells microplate. The different samples and solutions were then mixed in the following order: First 10 µL of the bovine insulin solution was transferred into the microplate containing the samples and the two solutions were mixed by aspirating/refouling 10 µL of the mixture 10 times. Then 15 µL of the resulting mixture was transferred into the plate containing the matrix and the two solutions were mixed by aspirating/refouling 10 µL of the mixture 10 times. 2 µL of the resulting sample/bovine insulin/matrix mixture was then immediately deposited as a drop on each spot of a MALDI MTP384 Polish steel target (Bruker Daltonics, Bremen, Germany). This was made possible by the creation and 3D printing of an adapter to allow the use of the MALDI target by the robot. Its design is presented in supplementary information (Figure S1). The drops were then dried at room temperature, and the MALDI target was introduced in the source chamber of an Autoflex Speed (Bruker Daltonics, Bremen, Germany). All MALDI-TOF analyses were performed in linear positive mode using an in-house method for insulin detection (LP_5-20_kDa.par), created from the manufacturer’s automatic method LP_5-20_kDa.par. Equipment parameters were as follow: voltage values of ion sources #1 and #2 set as 19.00 and 16.50 keV respectively; voltage values of reflectron #1 and #2 set as 21.00 and 9.50 keV respectively; lens tension 8.00 keV; pulsed extraction 120 ns; laser intensity between 60 and 70 %; laser Global attenuator Offset set to 41 %; Attenuator Offset set to 32 %, Attenuator Range set to 25 %; Detector Gain Voltage set to 2600 V (+300 V boost); Smartbeam Parameter set to ultra and Sample Rate and Digitizer Settings set to 4.00 GS/s. The MS signals were acquired by summing 5000 laser shots per spectrum. Prior to each analysis, the spectrometer was calibrated using the monoisotopic values of the manufacturer’s Protein Calibration Standard I calibrant (Bruker Daltonics, USA), containing Insulin ([M+H]^+^ - m/z=5734.51), Ubiquitin I ([M+H]^+^ - m/z=8565.76), Cytochrom C ([M+H]^+^ - m/z=12360.97), Myoglobin ([M+H]^+^ - m/z=16952.30), Cytochrom C ([M+2H]^2+^ - m/z=6180.99) and Myoglobin ([M+2H]^2+^ - m/z=8476.65). Calibrant was prepared by mixing 5 µL of calibrant diluted mixture prepared according to manufacturer’s specification with 5 µL of a 10 mg.mL^-1^ HCCA matrix in 50/49.9/0.1:acetonitrile/MS-grade water/trifluoroacetic acid, and 2 µL were then spotted on a Polished Steel 384 MALDI MTP target. Mass spectra of the sample were first visualized using FlexAnalysis software (version 3.4; Bruker Daltonics), after their calibration. To produce a comprehensive document, providing the intensity and area ratio of the murine and bovine insulin, as well quality control, in a way that is simple and easily understood by non-specialist experimenters, a short program was coded. It was written in VBA (code available in Supporting Information), and linked to an Excel sheet (Microsoft, Redmond, USA). After an automated export for the peak list from the mass spectra using FlexAnalysis software into a dedicated Excel sheet, this program can first perform an automated quality control, based on the intensity and the mass of the bovine insulin. Then it automatically detects of the murine insulin and performs the ratio in area or intensity compared to bovine insulin, depending on user’s preference. Finally, it sorts up all information and presents it in a clear colored view as a plate map, associated with ratio values and corresponding bar charts to quickly see the compounds that produce the highest signal compared to blank and others.

### Measurement of transfection efficiency by flow cytometry

To assess the transfection efficiency, Min6 cells were transfected with a siGLO (Green Transfection Indicator D-001630-01-05, Dharmacon) labelled with Fluorescein (CF/FAM/FITC)), using different concentrations of siGLO and transfection reagent. Transfection efficiency was determined 48 h after transfection using flow cytometry. Cells were acquired on a BD LSR Fortessa flow cytometer, and fluorescence of transfected min6 cells was quantified by flow cytometry.

### Immunofluorescence

Images from transfected Min-6 cells with siGLO and labeled with Hoechst 33342 (40 ng/mL, Thermofisher) were acquired with the microscope IN Cell Analyser 6000 (GE Healthcare Life Sciences) in 10X magnification in non-confocal mode with DAPI filter set (ex.405/em.455nm) and FITC filter set (ex.488/em.525nm) on the automated high content screening platform (Agilent Technologies, Santa Clara, USA) (Equipex Imaginex Biomed, Institut Pasteur de Lille, France).

### RNA extraction and quantitative real-time Polymerase Chain Reaction (qRT-PCR)

Total RNA was extracted from Min6 cells using the RNeasy Plus Microkit (Qiagen) following manufacturer’s instructions. Gene expression was measured after reverse transcription by quantitative real-time PCR (qRT-PCR) with FastStart SYBR Green master mix (Roche) using a LightCycler Nano or LC480 instruments (Roche). qRT-PCR results were normalized to endogenous cyclophilin reference mRNA levels. The results are expressed as the relative mRNA level of a specific gene expression using the formula 2^- ΔCt^. The complete list of primers is presented in Supplementary table 2.

### Immunoblotting experiments

Cells were washed with cold PBS and lysed in radioimmunoprotein-assay (RipA) buffer (10 mM Tris/HCl, 150 mM NaCl, 1 % (v/v) Triton X-100, 0.5 % (w/v) sodium deoxycholate, 0.1 % (w/v) sodium dodecyl sulfate and protease inhibitors, pH 7.4) and maintained under constant agitation for 30 min. Cell extracts were then centrifuged at 16,000 *g* for 20 min at 4 °C). Protein concentration was determined by the BCA protein assay kit according to the manufacturer’s instructions. Equal amounts of protein were resolved on 10 % SDS-PAGE under reducing conditions and proteins were transferred to a nitrocellulose membrane. Blots were incubated with primary antibodies directed against PDX1 (1:1000, Abcam, ab47267), alpha-tubulin (1:1000, Invitrogen, 32-2700), washed three times with PBS-0.05% tween and followed by incubation with secondary antibodies directed against goat anti-mouse or rabbit HRP conjugated (1:5000, Sigma-Aldrich). The visualization of immunoreactive bands was performed using the enhanced chemiluminescence plus western blotting detection system (GE Healthcare). Quantification of protein signal intensity was performed by volume densitometry using ImageJ 1.47t software (NIH).

### Statistical analysis

Data are expressed as mean ± SEM. Statistical analysis were performed using GraphPad Prism 9.3 software with two-way ANOVA with Bonferroni’s posttest for multiple comparisons as indicated in the figure legends. Differences were considered statistically significant at *P* value <0.05 (^*^<0.05; ^**^<0.01; ^***^<0.001; ^****^<0.0001). For the high-throughput screens, a score similar to the statistical Z-score for each test compound was calculated using the formula: Z *score =* (*X – µ*)*/ σ* where X is the murine insulin relative intensity from a compound-treated well, µ is the murine insulin relative intensity from the DMSO-treated wells on the same plate, and *σ* is the standard deviation of the murine insulin relative intensity signals of the DMSO-treated wells across all plates.

## Results

### Automation of Glucose□stimulated insulin secretion (GSIS) assay

In order to develop an automated protocol to measure insulin secretion in the mouse insulinoma Min6 cell line, we first miniaturized the GSIS protocol in 384-well plates (Supplementary Figure 2) and validate that these cells do respond to glucose stimulation in these culture conditions. Min6 cells were seeded and, 48 hours later, the cells were subjected to GSIS. For MALDI-TOF mass spectrometry analysis, removing BSA from the KRB buffer is a key step to limit BSA interferences and salt contaminations. Therefore, after one hour of starvation, the cells were washed 5 times with BSA-free KRB buffer supplemented with 2.8 mM glucose and incubated for 1 h at 37 °C. Then, half of the supernatant was collected using a Bravo liquid handler, and the cell plates were complemented with KRB buffer containing glucose at 20 mM and incubated for 1 h at 37 °C. The high glucose fractions were then collected and cells were lysed to measure insulin content.

To confirm the efficiency of the automated protocol, GSIS samples were subjected to an ELISA assay to measure insulin secretion in response to low and high glucose concentrations. Randomly selected samples from two separate 384-well plates were measured. Our data show that Min6 cells display a secretion rate which increases significantly after glucose stimulation, which represents 3 to 5 % of the total insulin content (Figure 1A) or a 6 to 8-fold increase of insulin secretion when Min6 cells were stimulated from 2.8 mM to 20 mM of glucose (Figure 1B). These data demonstrate that our GSIS protocol is functional in 384-well plates and that GSIS of Min6 cells can be fully automated in 384-well plates following our protocol.

**Figure 1.**
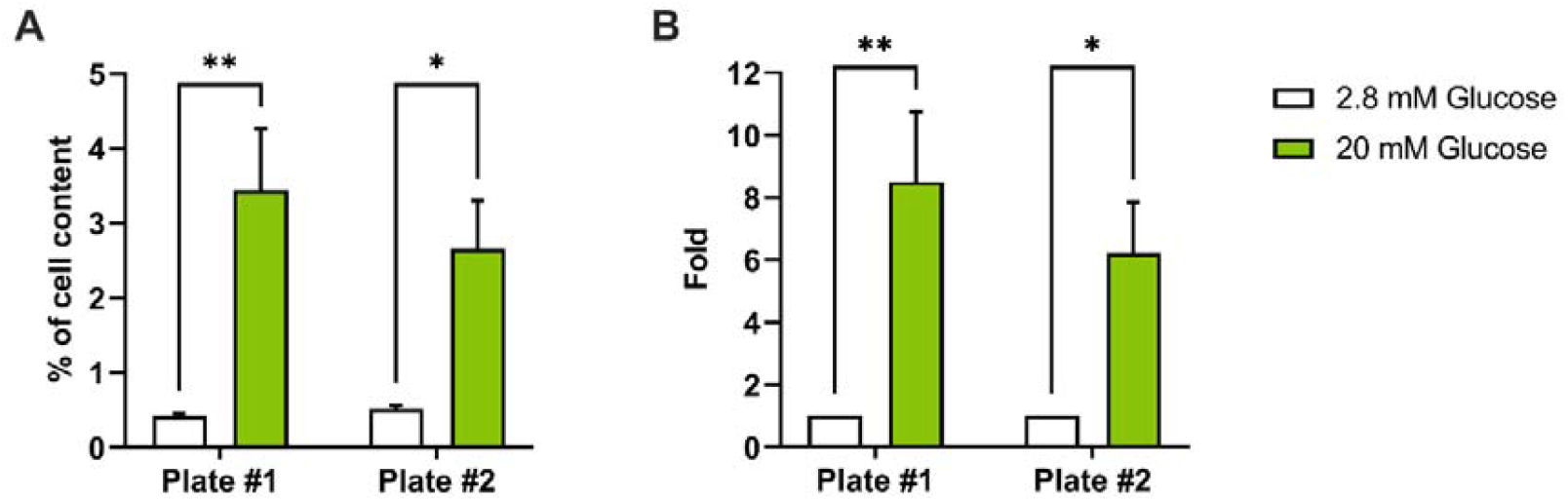
Automated GSIS is efficient to measure insulin secretion of Min6 cells. Min6 cells were cultured in 384-well plates and were subjected to automated GSIS. (A-B) Insulin concentration was measured using ELISA and GSIS results were expressed as percentage of insulin content (A, n □ =□ 5) or fold induction over 2.8 mM glucose (B, n □ = □ 5). Statistical analyses were performed using two-way ANOVA with Bonferroni post-test analyses. Results are presented as means ± SEM. * p<0.05 ; ** p<0.05.

### Automated analysis of GSIS in 384-well plates through MALDI-TOF mass spectrometry

Having established the miniaturized GSIS protocol, we next wanted to quantitatively detect insulin from GSIS assays through an automated approach that should be at least as sensitive, reliable and quantitative as ELISA and that can be automated. To this end, we selected MALDI-TOF mass spectrometry. First, we set up a spiked-based strategy that allows a precise quantification and normalization of the insulin protein present in our samples. Since the murine insulin has a molecular weight of 5803 Da, we selected bovine insulin as an internal standard, which has a molecular weight of 5733 Da and is assumed to present an ionizability very close to the one of murine insulin as its aminoacid composition is very similar. To quantitatively measure insulin from GSIS supernatants, samples were spiked with known concentrations of bovine insulin. Following MALDI-TOF mass spectrometry, a final mass spectrum was obtained, corresponding to the sum of 5,000 laser shots (Figure 2A). The relative intensity emitted by the detected murine insulin was normalized to the relative intensity of our spiked bovine insulin internal standard. Importantly, we observed that mixing our internal standard with our samples did not affect the signal strength of the detected murine insulin (Figure 2B). Then, to demonstrate the reliability of MALDI-TOF mass spectrometry to detect insulin, samples from GSIS experiments performed in 384-well plates were analyzed through ELISA assay and data were compared to MALDI-TOF mass spectrometry results. When we measured automated GSIS through ELISA or MADI-TOF mass spectrometry, we could not observe differences in GSIS results between these two approaches. The analysis of two independent plates demonstrated that Min6 cells secreted between 6 to 8 times more insulin at 20 mM glucose compared to 2.8 mM glucose, independently of the method of detection, *i*.*e*. mass spec or ELISA (Figure 2C). These results demonstrate a strong reliability between both detection methods to measure insulin protein and suggest that MALDI-TOF mass spectrometry is as sensitive, as reliable and as quantitative as ELISA for GSIS measurements.

**Figure 2.**
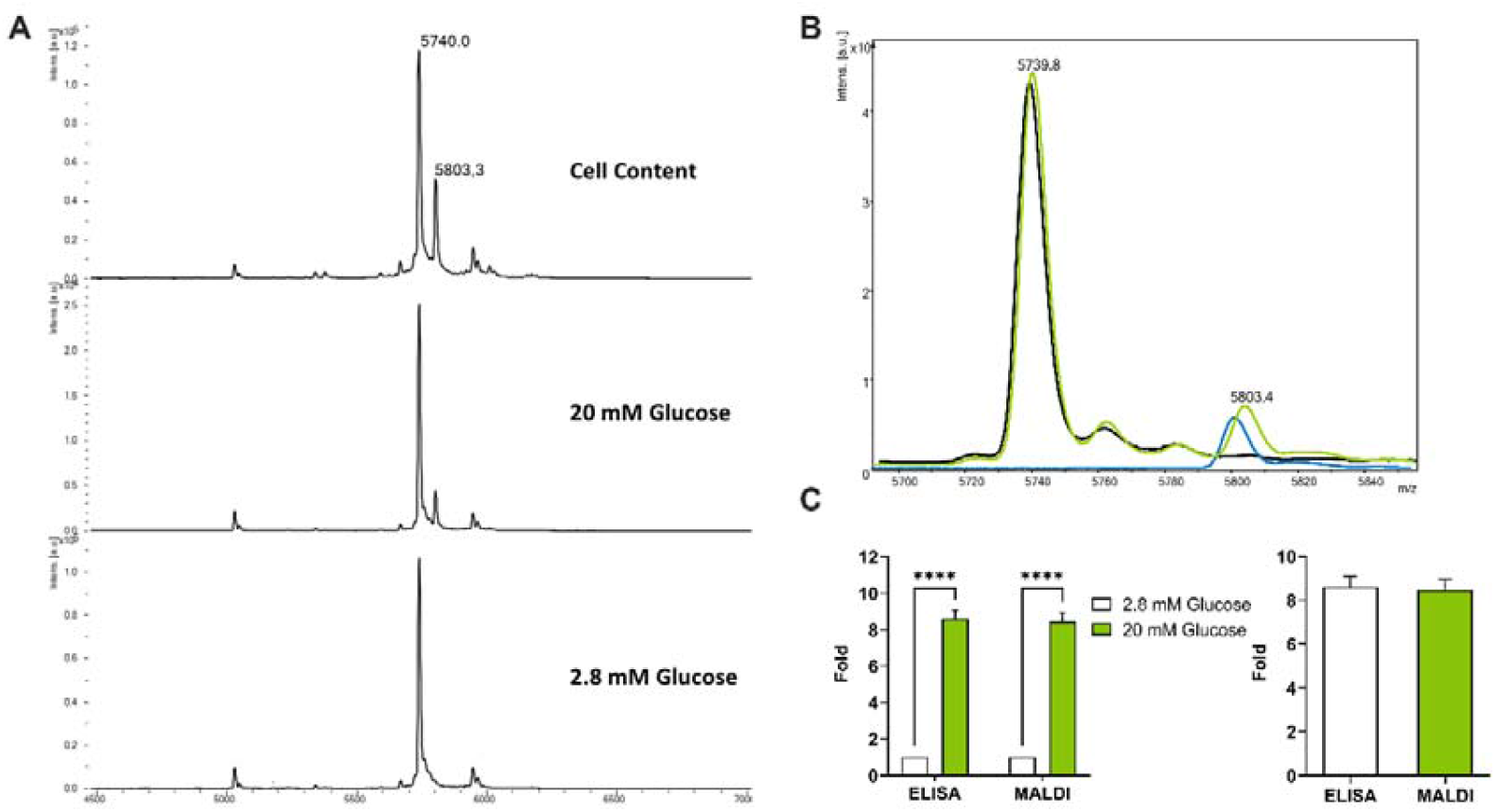
Measurement of insulin secretion from Min6 cells through MALDI-TOF mass spectrometry. (A) Mass spectra of the intracellular insulin content, 20 mM and 2.8 mM of glucose secretomes (B) Superposition of the different mass spectra of murine and bovine insulin (black: bovine, blue: murine, green: mixture) (C) Quantification of insulin secretion after stimulation with glucose by MALDI-TOF mass spectrometry. Min6 cells were cultured in a 384-well plate and then subjected to automated GSIS and insulin secretion was measured comparing ELISA (n□=□4) and MALDI-TOF (n=16) as detection methods. Samples from two independent plates were analyzed. Statistical analyses were performed using two-way ANOVA with Bonferroni post-test analyses. Results are presented as means ± SEM. * p<0.05 ; ** p<0.01 ; *** p<0.001 ; **** p<0.0001

### Automated siRNA transfection combined with GSIS

We next wanted to apply the automated GSIS protocol to the discovery of new potential genes involved in insulin secretion. Following our experimental strategy described above, a fully automated siRNA reverse transfection protocol was implemented with the aim to detect variations of insulin secretion upon knocking-down specific genes. We first determined the optimal transfection conditions using a 6-FAM fluorescent-labeled control siRNA. This step is crucial to ensure an efficient transfection rate to potentially obtain a significant knock-down of the expression of target genes. Using siGLO as fluorescent control to assess transfection efficiency, we could determine the optimal concentration for transfection reagents, siRNA and time of incubation (Supplementary Figure 3). Once the automated miniaturized transfection and GSIS protocols were validated, we tested two positive control siRNA *Pdx1* and *Kcnj11*, two genes previously shown to negatively modulate insulin secretion. Knock-down of *Pdx1* gene in Min6 cells was validated both at the transcriptomic and protein levels (Supplementary Figure 4). Upon knock-down, GSIS samples were then analyzed by MALDI spectrometry and we detected lower relative intensities for murine insulin in cells treated with siRNA targeting *Pdx1* and *Kcnj11* (Figure 3A and 3B). Indeed, intensity ratios under 2.8 mM and 20 mM glucose conditions showed a 10-fold increase of insulin secretion for siControl-treated Min6 cells, whereas a 6-fold increase in si*Pdx1* -treated cells and a 4-fold increase in si*Kcnj11*-treated cells were observed (Figure 3C). These results were further independently confirmed through ELISA assay where siRNAs targeting *Kcnj11* and *Pdx1* induced a significant decrease in insulin secretion after a stimulation with 20 mM glucose (Figure 3D). Again, mass spectrometry analysis further confirmed the results obtained through ELISA approaches (Figure 3E). Altogether, these data demonstrate that an automated siRNA reverse transfection protocol is efficient to evaluate potential functional effects of knocking-down genes in Min6 cells and suggest that this approach provides a robust automated platform to evaluate GSIS in HTS approaches.

**Figure 3.**
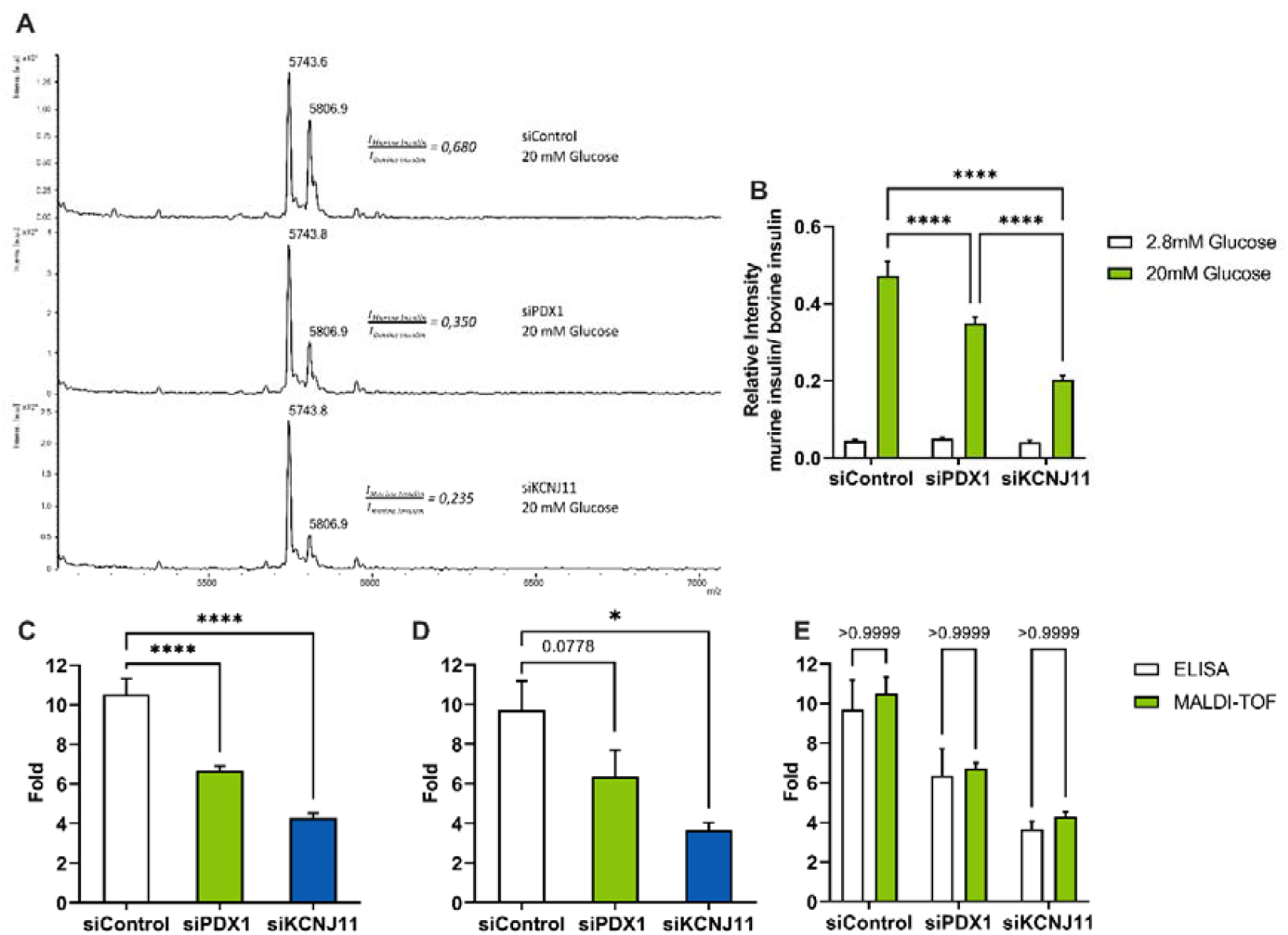
Automated GSIS of siRNA-transfected Min6 cells analyzed through MALDI-TOF mass spectrometry. (A) Mass spectra of the 20 mM secretomes of Min6 cells transfected with siControl, si*Pdx1* and si*Kcnj11* (B) Quantification of mass spectra of Min6 cells transfected with siControl, si*Pdx1* and si*Kcnj11*subjected to GSIS, using relative intensities of murine and bovine insulin spectrum area. (C, D) Glucose stimulation effect on insulin secretion presented as the fold of insulin secretion in 20 mM glucose over 2.8 mM Glucose. Insulin levels were measured from Min6 cells transfected by siControl, si*Pdx1* and si*Kcnj11* through MALDI-TOF mass spectrometry (C, n=16) or ELISA (D, n=4). (E) Comparison of quantification results obtained by mass spectrometry and ELISA assays. Statistical analyses were performed using two-way ANOVA with Bonferroni posttest analyses. Results are presented as means ± SEM. * p<0.05 ; ** p<0.01 ; *** p<0.001 ; **** p<0.0001

### Automated chemical treatment combined with GSIS

As a proof of concept, we next wanted to implement HTS strategies using our Min6 automated GSIS protocol coupled to mass spectrometry analysis. To demonstrate the feasibility of our approach, the use of pharmacological activators (Repaglinide) and inhibitors (Diazoxide) of insulin secretion has been undertaken to observe insulin secretion variations in Min6 cells. After treatment of the cells with these compounds, MALDI-TOF mass spectrometry analysis was employed to measure insulin secretion in response to these drugs. Our data revealed that these compounds were effective on modulating glucose-stimulated insulin secretion (Figure 4A), as demonstrated using automated MALDI TOF mass spectrometry or ELISA assays. Indeed, upon stimulation with 2.8 mM glucose condition, Min6 cells treated with 100 nM of repaglinide had a higher basal secretion rate than untreated cells or cells incubated with 100 μM of diazoxide (Figure 4B). Upon 20 mM glucose condition treatment, a 2-fold increase and a 2-fold decrease in insulin secretion in the presence of repaglinide and diazoxide, respectively, was observed (Figures 4C and 4D), which were again confirmed through ELISA assays (Figure 4E). In addition, the intracellular insulin content of Min6 cells exposed to repaglinide was decreased by 30 % when compared to other conditions (Supplementary Figures 5A, B and C), yet the total amount of insulin was similar in Min6 for all conditions of treatments (Supplementary Figures 5D).

**Figure 4.**
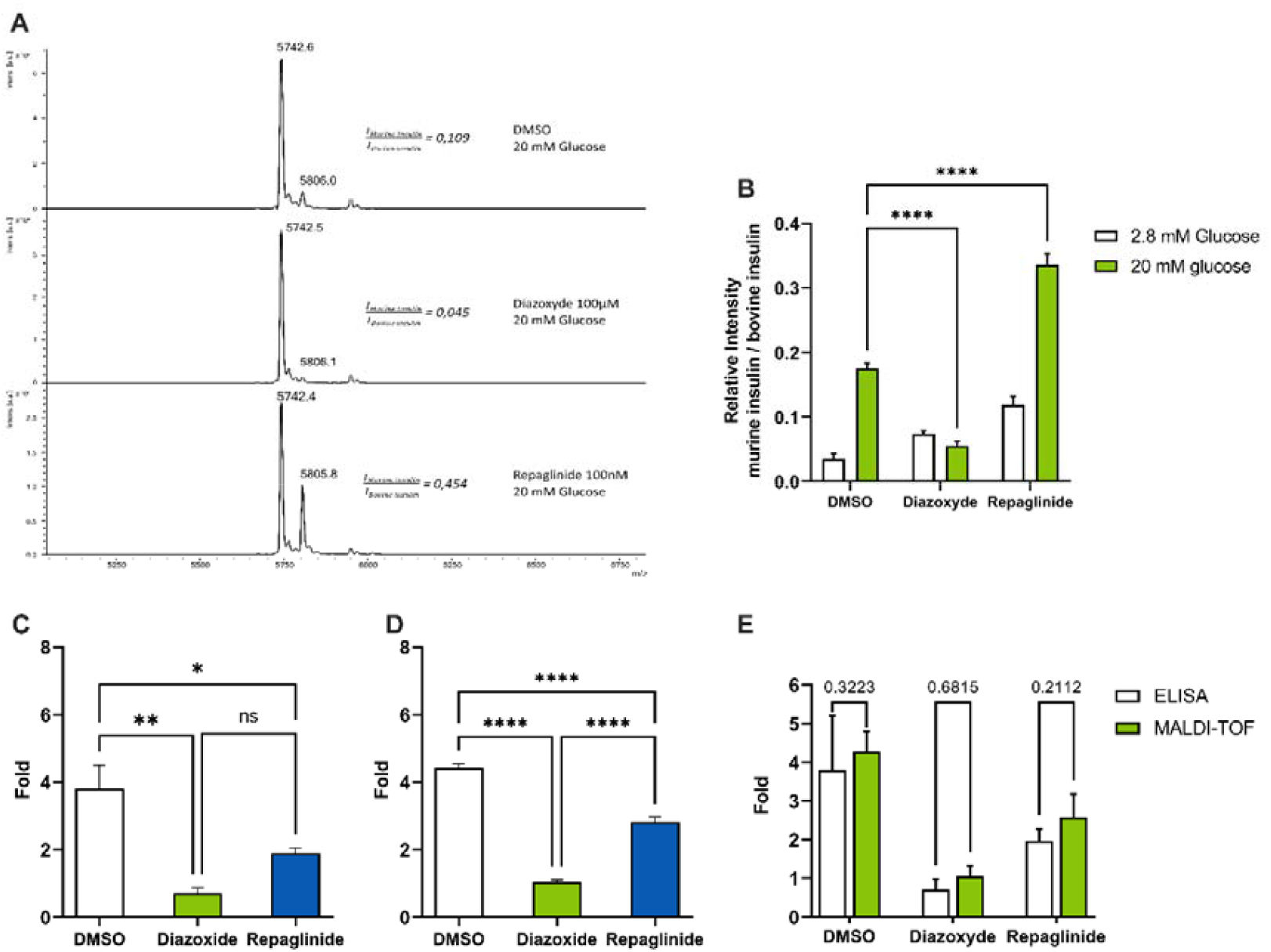
Min6 insulin secretion analysis of Min6 cells treated with diazoxide and repaglinide by MALDI-TOF mass spectrometry. (A) Mass spectra of the 20 mM secretomes of Min6 cells treated with DMSO, diazoxide 100 µM and repaglinide 100 nM Quantification of insulin secretion after stimulation with glucose 2,8 mM and 20 mM of Min6 cells treated with DMSO, diazoxide 100µM and repaglinide 100 nM (B,D) by MALDI-TOF mass spectrometry and (C) by ELISA. (E) Comparison of results obtained by mass spectrometry and ELISA assay. Two-way ANOVA with Bonferroni posttest analyses. Error bars are ± SEM and n□=□4 for ELISA, n=32 for MS. * p<0.05 ; ** p<0.01 ; *** p<0.001 ; **** p<0.0001

As expected, cells stimulated with repaglinide showed decreased intracellular insulin content when compared to control, untreated cells. Indeed, treating Min6 cells with repaglinide induced a 70 % decrease of the intracellular insulin content, and, conversely, diazoxide increased by 30 % insulin content compared to control, untreated cells (Figure 5A). Thanks to our high throughput MALDI-TOF approach, we analyzed 160 vehicle-treated samples, 62 repaglinide-treated samples and 62 diazoxide-treated samples and observed that their analysis followed a normal distribution (Figure 5B and Supplementary Figure 6). In addition, we could repeat this experiment with other bioactives molecules known to induce insulin secretion, such as forskolin or IBMX, and confirmed our results obtained with repaglinide (Supplementary Figure 7). Indeed, after GSIS, the intensity levels corresponding to insulin observed for the cells treated with different secretagogues were markedly lower than the control, untreated cells, suggesting that these compounds may provoke intracellular insulin leakage (Supplementary Figure 7).

**Figure 5.**
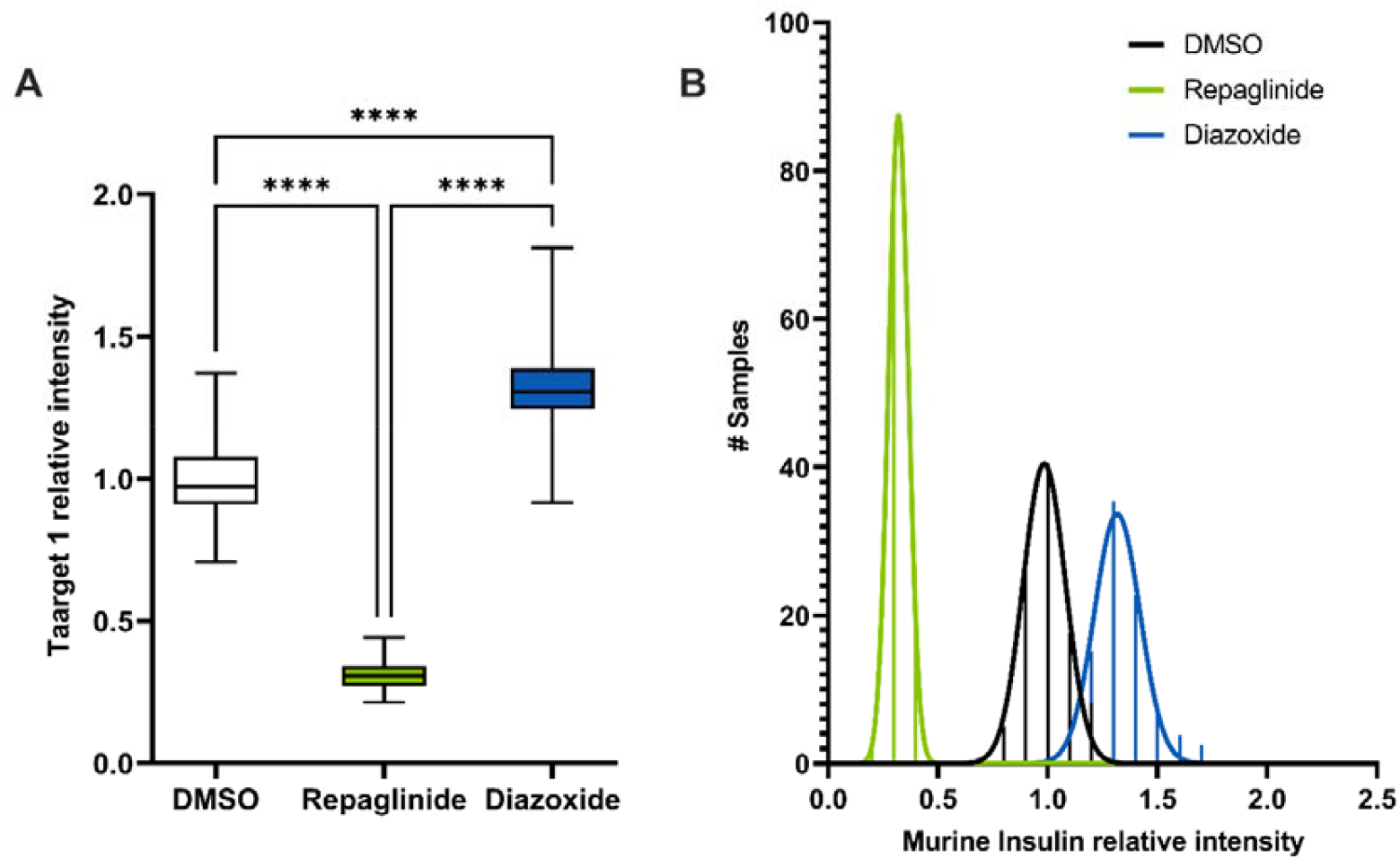
Analysis of insulin content of Min6 cells treated with diazoxide and repaglinide through MALDI-TOF mass spectrometry. Quantification of intracellular insulin content of Min6 cells treated with DMSO, 100µM diazoxide and 100nM repaglinide through MALDI-TOF mass spectrometry. Statistical analyses were performed using two-way ANOVA with Bonferroni posttest analyses. Results are presented as means ± SEM. n□=□4 for ELISA, n=160 for DMSO and n=80 for Repaglinide and Diazoxide. * p<0.05 ; ** p<0.01 ; *** p<0.001 ; **** p<0.0001.

### Automated library screening to mesure insulin intracellular content after GSIS

Since our automated cell culture protocol, MALDI-TOF mediated insulin detection and quantification and GSIS pipeline is adapted for high-throughput applications, we performed a pilot screening experiment in Min6 cells using a collection of 1,600 compounds from ASINEX library and DMSO, repaglinide and diazoxide as controls. Here, we focused on measuring insulin content only. As observed previously, repaglinide induced a significantly lower intracellular insulin content compared to the negative controls (Figure 6).

**Figure 6.**
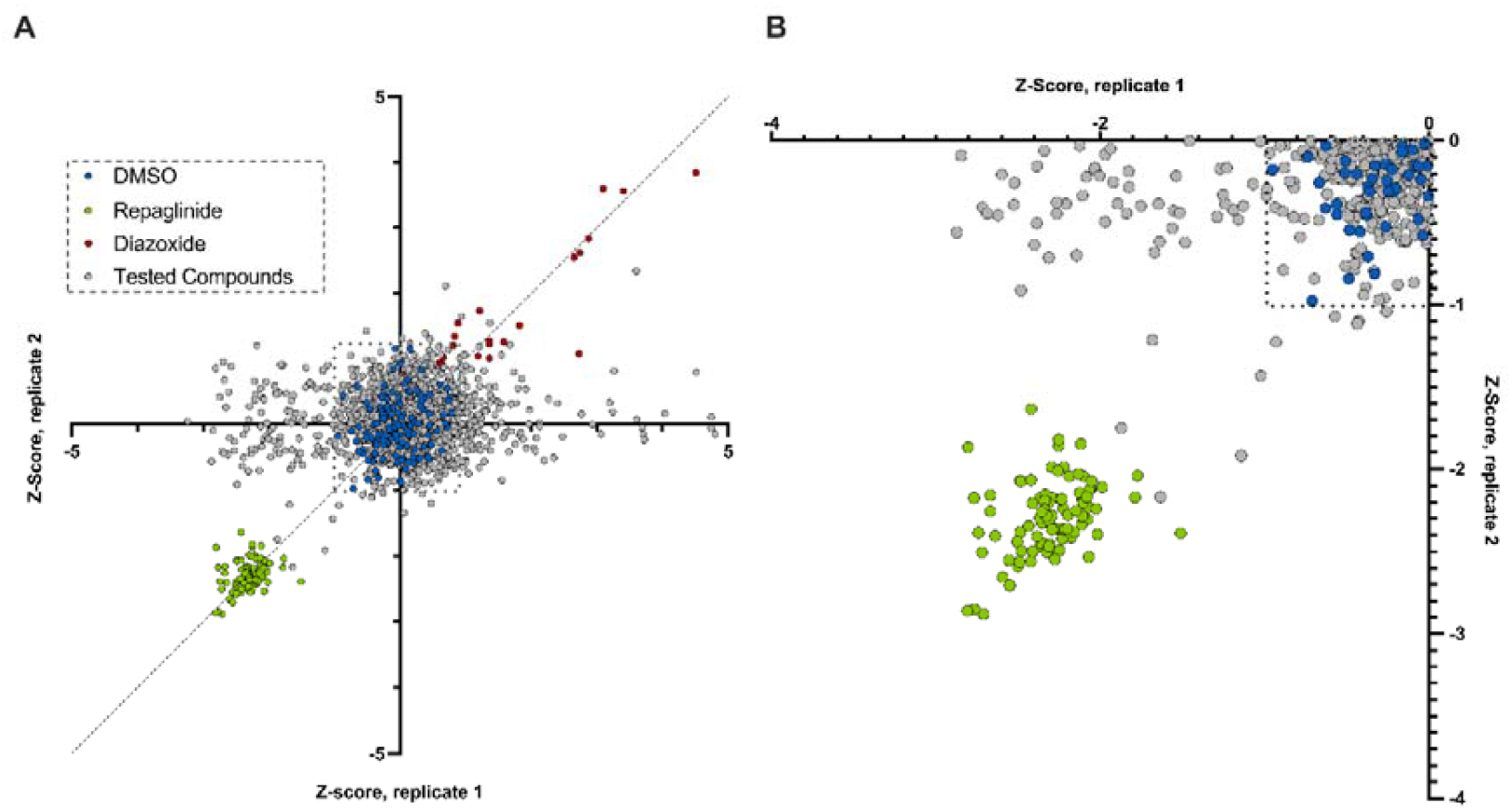
High-Throughput chemical screen of Min6 cells identifies small molecules that amplify insulin release and decrease insulin intracellular content after glucose stimulation. Min6 cells were treated with 1,600 chemical compounds and insulin content was measured using MADI-TOF mass spectrometry. Compounds from the chemical library are in grey, and molecules that potentially have an effect on insulin content are indicated by a grey arrow. Repaglinide (green) was used as a positive control, whereas diazoxide (red) was used as an inhibitor of insulin secretion.

This screening was carried out in duplicate in order to be able to calculate a z-score for all the compounds tested. The DMSO control cells presented a z-score comprised in a window between z = -1 and z = 1. The cells treated with repaglinide had a z-score between z = - 1.5 and z = -3. We assumed that cells treated with compounds with the same order of magnitude of repaglinide z-score may be considered as molecules that stimulate insulin secretion. Conversely, compounds leading to a z-score close to controls or diazoxide may be considered as, at least, non-effective or inhibitors of insulin secretion. Among the 1600 compounds tested, we observed that some molecules decreased insulin content after glucose stimulation (Figure 6), suggesting that they may stimulate insulin secretion.

Amongst those molecules, we observed different compounds with z-score close to those of repaglinide, the positive control. Altogether, our results demonstrate that our approach using MALDI-TOF mass spectrometry can be automated and efficiently used for HTS of libraries of siRNA or chemical compounds.

## Discussion

Here, we describe a new automated methodology to measure accurately insulin secretion in vitro and identify potential modulators of insulin secretion in pancreatic beta cells. This new process, based on automated GSIS and MALDI-TOF mass spectrometry, was developed in a 384-well plate format to enable high-throughput analyses and identify new chemical substances or genes that potentially modulate insulin secretion or cell content. We demonstrated the efficiency of our assay by performing a pilot experiment using 1,600 chemical compounds and selected siRNAs, in the mouse Min6 beta cell line. Indeed, we have developed and implemented this technological process to test the efficiency of screening different chemical compounds, but also siRNA targeting bona fide β-cell identity genes. Unlike other techniques such as ELISA as-say, our protocol is based on direct insulin detection and not on antigen recognition. In addition, we demonstrate that the MALDI-TOF mass spectrometry strategy is as reliable as ELISA detection, while being much less expensive than existing insulin detection techniques.

High-throughput screening for insulin secretion is an approach with high potential for valorization. In the case of the insulin-secreting pancreatic β-cell, these approaches are limited by several technological constraints inherent to the methods for detection and quantification of this hormone (i.e., ELISA). To overcome these experimental limitations, several laboratories have reported the use of modified cell-based tools to indirectly measure the level of secretory activity of mouse-derived Min6 cells by fluorescence microscopy or measurement of modified peptide-C luciferase [6–9]. Although simple to manipulate, these artificial, genetically-engineered tools only very partially interrogate β-cell functionality as it assesses neither endogenous insulin pro-duction nor its secretion under glucose stimulation conditions. It is with the objective of addressing these two analytical criteria that we have proposed to deploy our complete procedure of automation of the GSIS protocol followed by MALDI-TOF mass spectrometry to envision, in the future, a siRNA library (approx. 20,000 genes) and chemical library (>100,000 compounds) screening strategy in the Min6 pancreatic β-cell model. A similar study has already been undertaken on the INS-1 832/13 pancreatic beta cell model, with the screening of 1200 compounds, but unlike our study, the compounds used have already known targets [10,11].

The scope of the technology is likely very broad, as MS detection is extremely versatile. The most direct applications being the high-throughput screening for other molecules that modulate the secretion of other peptide hormones such as, Glucagon-like peptide 1 (Glp-1)…) In the context of the development of personalized medicine, our analysis process could be deployed to define the most appropriate treatment to restore the hormonal secretion of a patient suffering from T2D. This concept of searching for alternative therapeutics could naturally be extended to other disciplinary fields such as oncology or the study of neurodegenerative diseases. Nevertheless, the use of our process in a precise context remains subject to the prior knowledge of the protein(s) of interest.

Although encouraging, our study has several limitations. The use of the murine β-cell model Min6 is convenient, but translation to human cells remains to be done. It may be relevant to confirm some of the target and/or small molecules identified in Min6 cells in the EndoC-BH1 human model, which has been shown to be useful for the identification of modulators of human beta-cell insulin secretion [12]. In addition, siRNA mediated knock-down may not be efficient enough to identify genes potentially involved in β-cell function. The use of CrispR/Cas9, as performed in human islets [13] or in EndoC-BH1 [14] may help to better identify genes that are directly involved in insulin secretion, strengthening the translation to human.

## Supplementary Materials

Table S1. List of siRNA used for Min6 cells transfection; Table S2. List of oligonucleotides used in qPCR experiments; Table S3. Relevant compounds identified as positive modulators of insulin secretion; Figure S1: 3D Printing adapter to allow the use of the MALDI target by Biomek NXp liquid handler; Figure S2: Schematic Representation of the GSIS pipeline; Figure S3: Efficiency tests for automated siRNA approach; Figure S4: Efficiency of automated transfection of Min6 cells; Figure S5: Min6 insulin secretion analysis of Min6 cells treated with diazoxide and repaglinide by MALDI-TOF mass spectrometry; Figure S6: Normal distribution of analysis of intracellular insulin content from Min6 cells treated with DMSO, repaglinide and diazoxide; Figure S7: High-throughput analysis of intracellular insulin content of min6 cells treated with DMSO, IBMX, diazoxide, repaglinide and forskolin after glucose-stimulated insulin secretion.

## Author Contributions

Conceptualization, C.P.D., E.H., A.H., X.G., F.O., and J.S.A.; Methodology, C.P.D., E.H., A.H., F.O., M.C., X.G. and J.S.A.; Software, E.H.; Validation, C.P.D., E.H., A.H., and J.S.A.; Formal analysis, C.P.D., and E.H.; Investigation, C.P.D., E.H., A.H., F.O., B.T., M.C., X.G. and L.R.; Resources, P.F., B.D., and S.P.; Data curation, C.P.D., E.H., and A.H.; Writing – original draft, C.P.D., and J.S.A.; Writing – Review & Editing, E.H., A.H and B.D.; Supervision, J.S.A.; Project administration, B.D., S.P., and J.S.A.; Funding acquisition, J.S.A.

## Acknowledgments

The authors thank the members of the INSERM U1283-EGID and INSERM UMR1167-RID-AGE for helpful discussions. The authors are grateful to Yannick Campion (SATT-Nord) for his support and his administrative contribution during the time of the project. Part of the automated workflow described here was set up and performed on the REALCAT platform, funded by a French governmental subsidy managed by the French National Research Agency (ANR) within the framework of the “Future Investments’ program (ANR-11-EQPX-0037). The Hauts-de-France region, FEDER, Ecole Centrale de Lille, and Centrale Initiatives Foundation are also warmly acknowledged for their financial contributions to the acquisition of the REALCAT platform equipment.

## Supplementary Data

**Figure S1:**
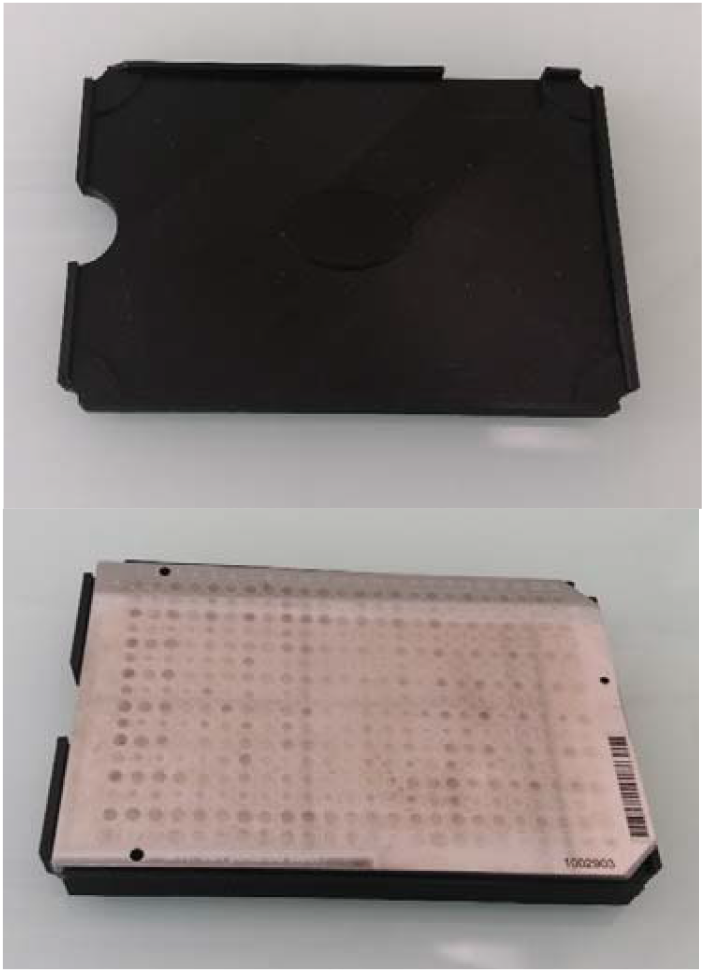
3D Printing adapter to allow the use of the MALDI target by Biomek NX^p^ liquid handler.

**Figure S2:**
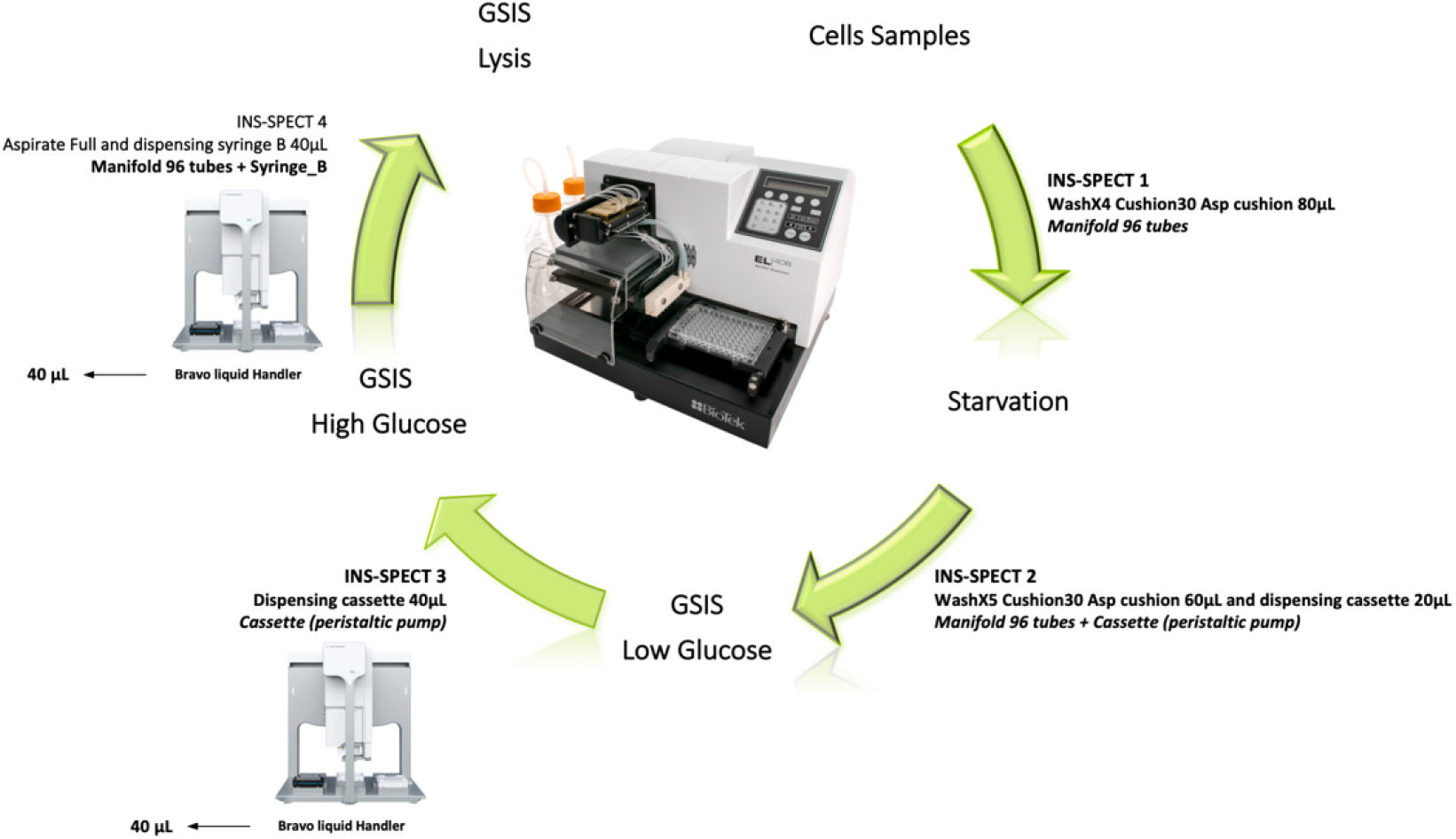
Schematic Representation of the GSIS pipeline.

**Figure S3:**
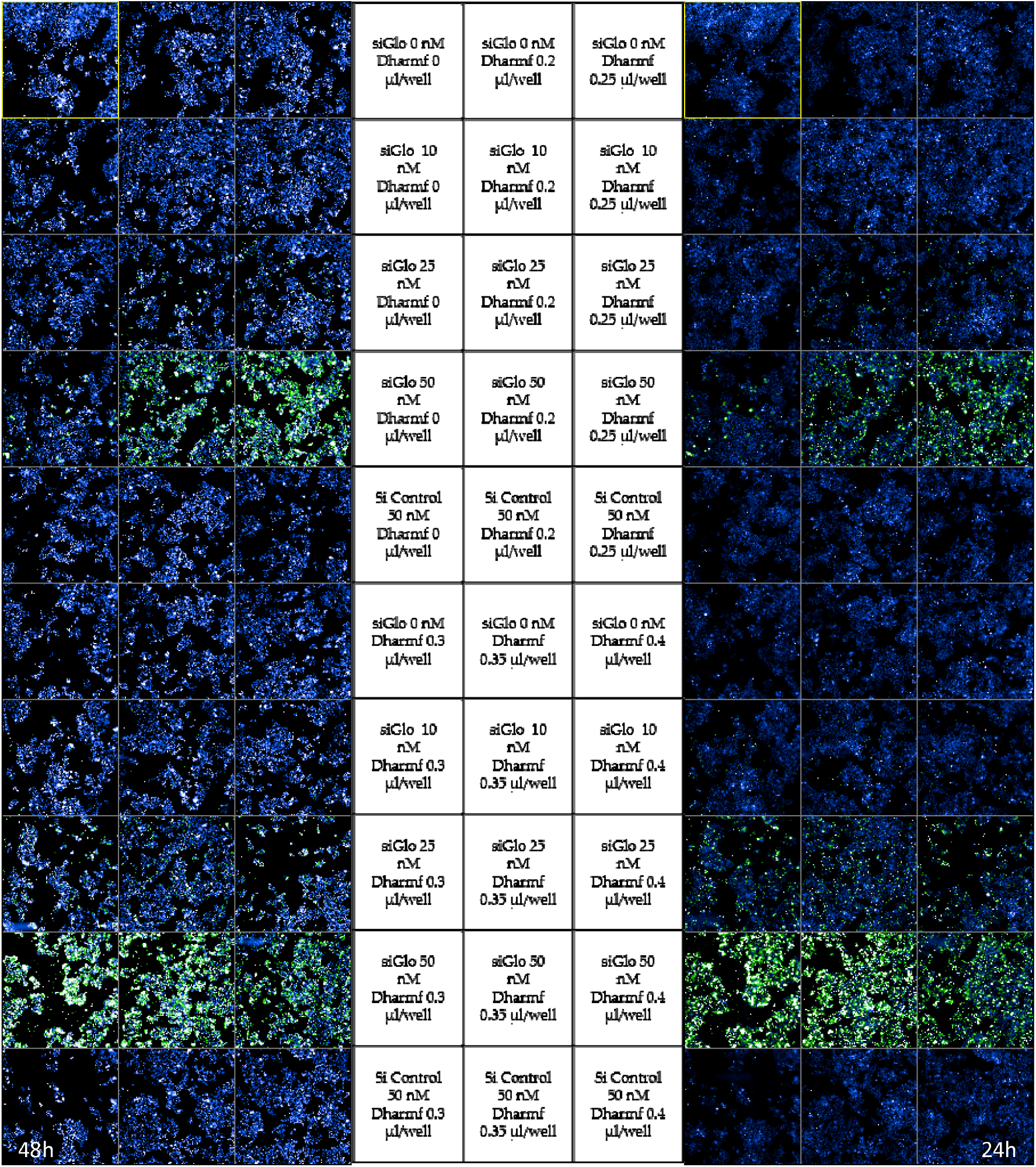
Efficiency tests for automated siRNA aproach. Confocal fluorescence microscopy images of cells transfected with siGLO and non-fluorescent siControl at different concentrations, time of incubation. Different concentrations for the transfcetant reagents is also shown.

**Figure S4:**
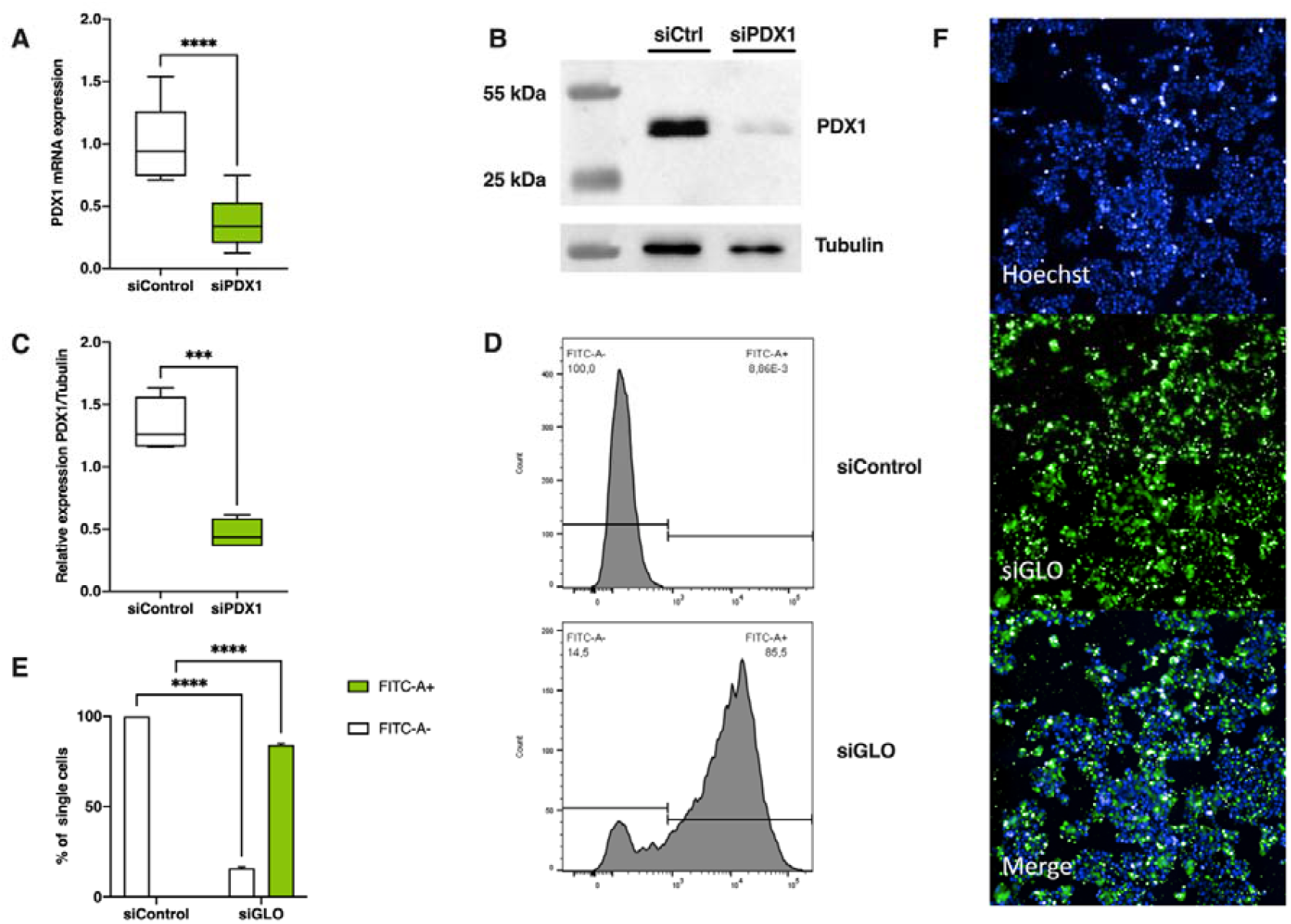
Efficiency of automated transfection of Min6 cells. (A) PDX1 mRNA expression from siCtrl and siPDX1 Min6 cells after after automated reverse transfection; (B, C) Western blot showing mouse PDX1 protein level in Min6 cells after automated reverse transfection with siRNA control and siPDX1; (D) Histogram and (E) graphical representation of the number of FITC-A- and FITC-A + cells after transfection with siGLO. (F) Confocal fluorescence microscopy images of cells transfected with siGLO. Two-way ANOVA with Bonferroni posttest analyses. Error bars are ± SEM and n□=□3.* p<0.05 ; ** p<0.01 ; *** p<0.001 ; **** p<0.0001.

**Figure S5:**
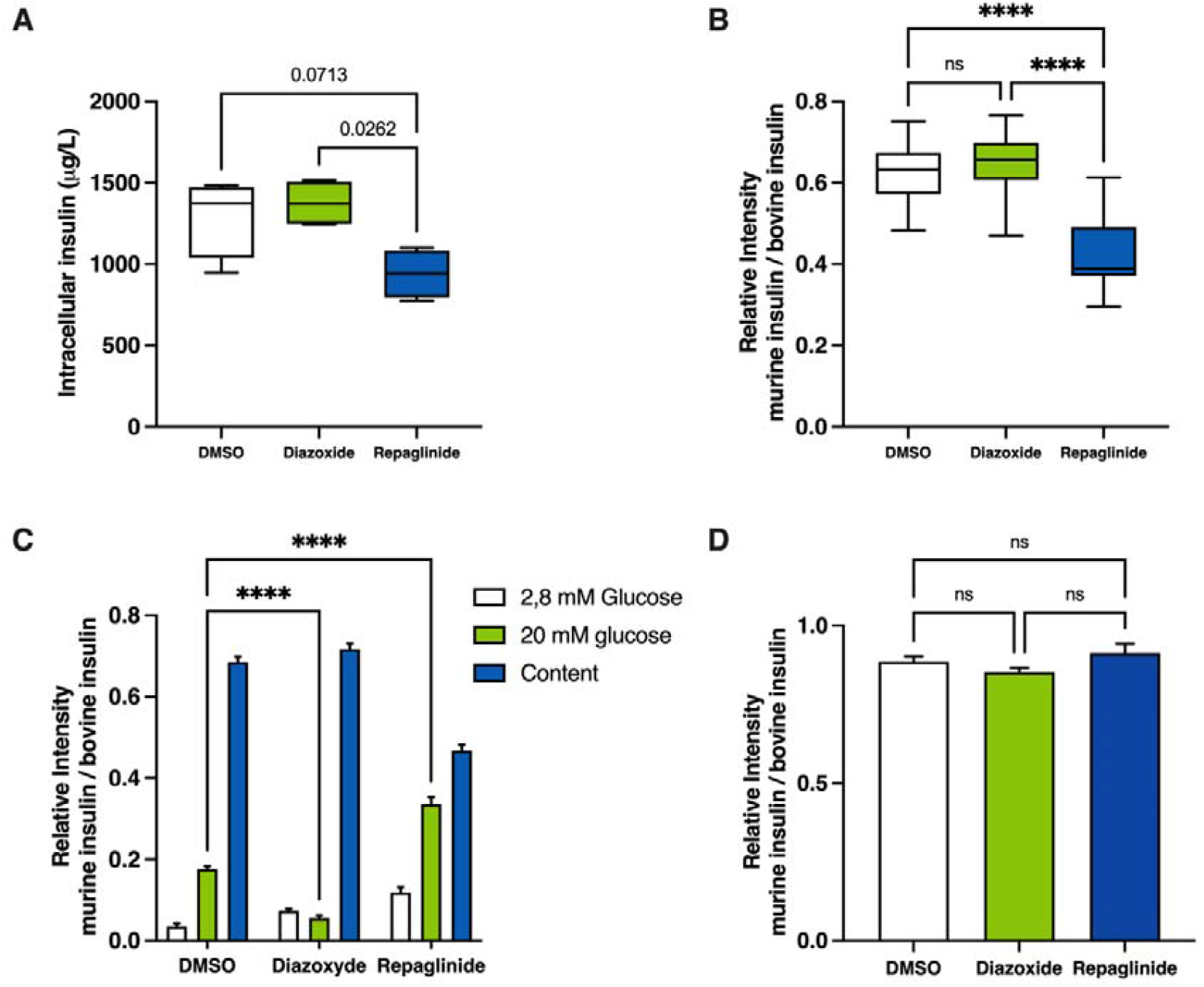
Min6 insulin secretion analysis of Min6 cells treated with diazoxide and repaglinide by MALDI-TOF mass spectrometry. Quantification of insulin intracellular content of Min6 cells treated with DMSO, diazoxide 100µM and repaglinide 100nM by MALDI-TOF mass spectrometry (A,C) and ELISA assay (B). Total insulin stored and secreted in min6 cells treated with DMSO, diazoxide 100µM and repaglinide 100nM by MALDI-TOF mass spectrometry. Two-way ANOVA with Bonferroni posttest analyses. Error bars are ± SEM and n□=□4 for ELISA, n=32 for MS. * p<0.05 ; ** p<0.01 ; *** p<0.001 ; **** p<0.0001

**Figure S6:**
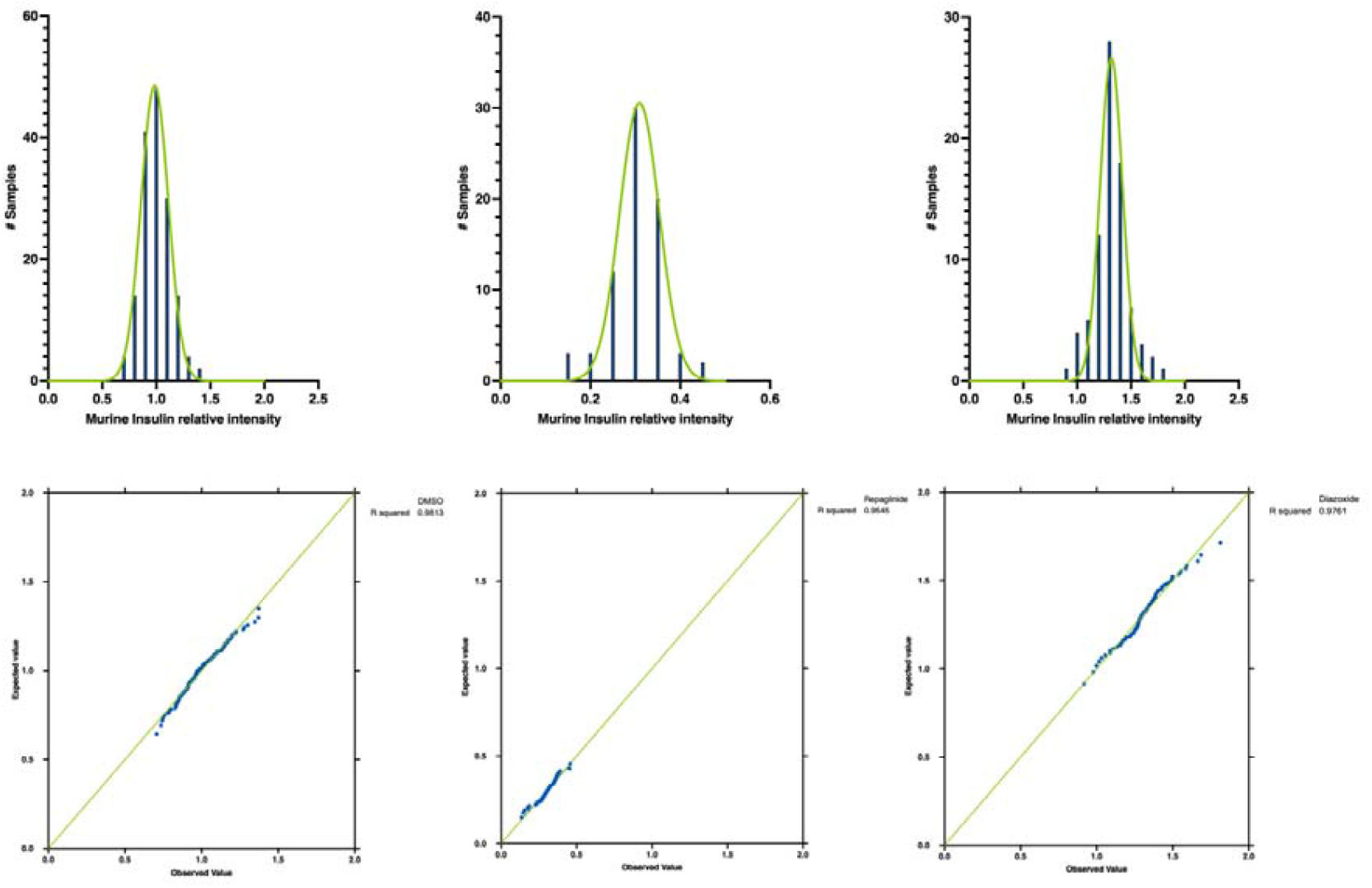
Normal distribution of analysis of intracellular insulin content from Min6 cells treated with DMSO, repaglinide and diazoxide.

**Figure S7:**
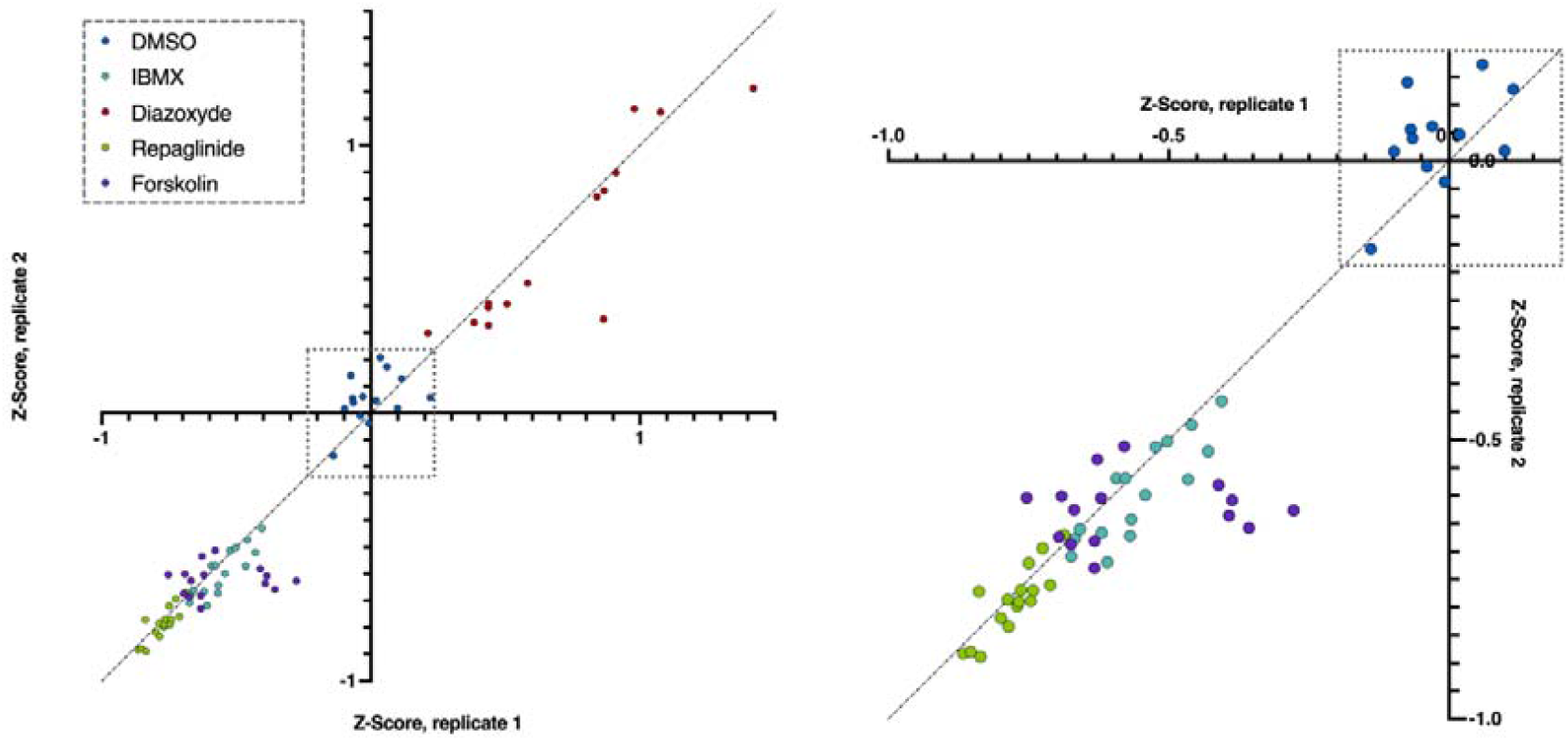
High-throughput analysis of intracellular insulin content of min6 cells treated with DMSO, IBMX, diazoxide, repaglinide and forskolin after glucose-stimulated insulin secretion.

**Table S1.**
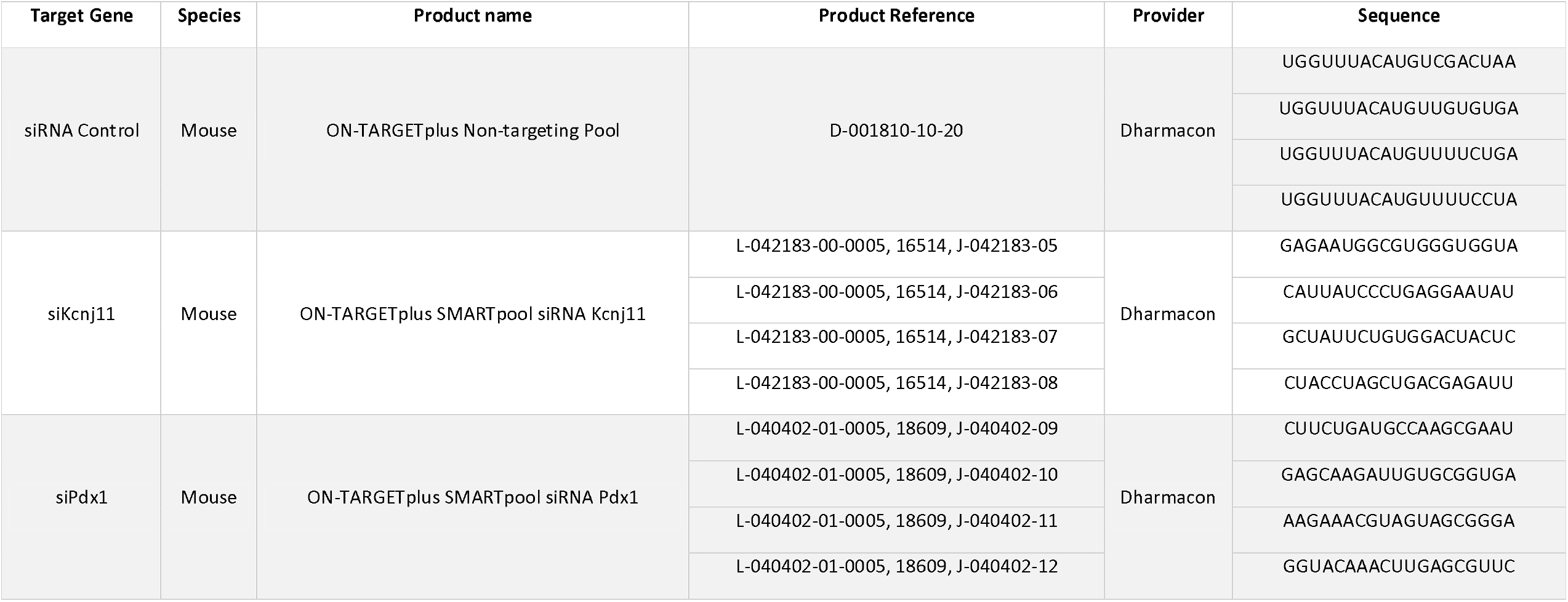
List of siRNA used for Min6 cells transfection

**Table S2.**
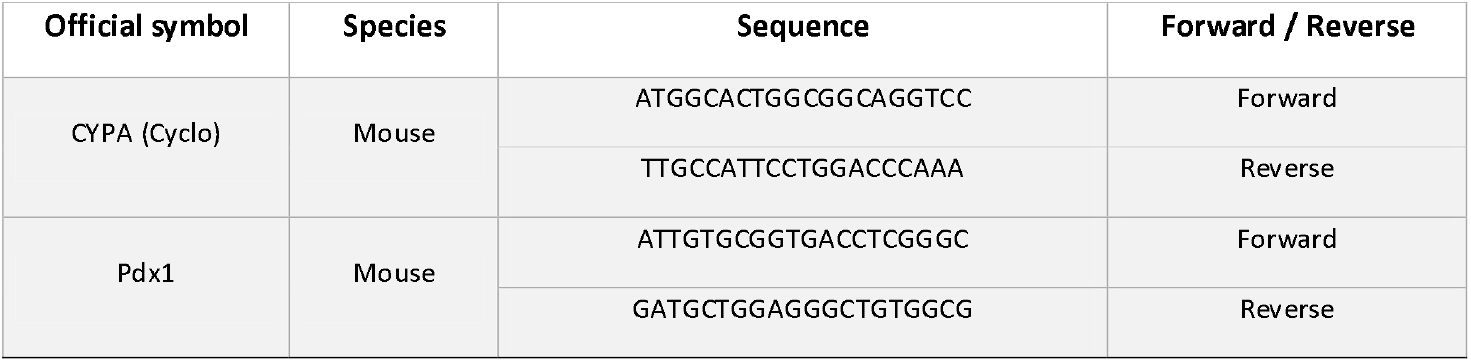
List of oligonucleotides used in qPCR experiments

## References

1. Lin, X.; Xu, Y.; Pan, X.; Xu, J.; Ding, Y.; Sun, X.; Song, X.; Ren, Y.; Shan, P.-F. Global, Regional, and National Burden and Trend of Diabetes in 195 Countries and Territories: An Analysis from 1990 to 2025. Sci Rep 2020, 10, 14790, doi:10.1038/s41598-020-71908-9.

2. Artasensi, A.; Pedretti, A.; Vistoli, G.; Fumagalli, L. Type 2 Diabetes Mellitus: A Review of Multi-Target Drugs. Molecules 2020, 25, 1987, doi:10.3390/molecules25081987.

3. Nisal, K.; Kela, R.; Khunti, K.; Davies, M.J. Comparison of Efficacy between Incretin-Based Therapies for Type 2 Diabetes Mellitus. BMC Medicine 2012, 10, 152, doi:10.1186/1741-7015-10-152.

4. Aguayo-Mazzucato, C. Functional Changes in Beta Cells during Ageing and Senescence. Diabetologia 2020, 63, 2022–2029, doi:10.1007/s00125-020-05185-6.

5. Heyduk, E.; Moxley, M.M.; Salvatori, A.; Corbett, J.A.; Heyduk, T. Homogeneous Insulin and C-Peptide Sensors for Rapid Assessment of Insulin and C-Peptide Secretion by the Islets. Diabetes 2010, 59, 2360–2365, doi:10.2337/db10-0088.

6. Kalwat, M.A.; Wichaidit, C.; Nava Garcia, A.Y.; McCoy, M.K.; McGlynn, K.; Hwang, I.H.; MacMillan, J.B.; Posner, B.A.; Cobb, M.H. Insulin Promoter-Driven Gaussia Luciferase-Based Insulin Secretion Biosensor Assay for Discovery of β-Cell Glucose-Sensing Pathways. ACS Sens 2016, 1, 1208–1212, doi:10.1021/acssensors.6b00433.

7. Burns, S.M.; Vetere, A.; Walpita, D.; Dančík, V.; Khodier, C.; Perez, J.; Clemons, P.A.; Wagner, B.K.; Altshuler, D. High-Throughput Luminescent Reporter of Insulin Secretion for Discovering Regulators of Pancreatic Beta-Cell Function. Cell Metabolism 2015, 21, 126–137, doi:10.1016/j.cmet.2014.12.010.

8. Pappalardo, Z.; Chopra, D.G.; Hennings, T.G.; Richards, H.; Choe, J.; Yang, K.; Baeyens, L.; Ang, K.; Chen, S.; Arkin, M.; et al. A Whole-Genome RNA Interference Screen Reveals a Role for Spry2 in Insulin Transcription and the Unfolded Protein Response. Diabetes 2017, 66, 1703–1712, doi:10.2337/db16-0962.

9. Kalwat, M.A. High-Throughput Screening for Insulin Secretion Modulators. In Exocytosis and Endocytosis: Methods and Protocols; Niedergang, F., Vitale, N., Gasman, S., Eds.; Methods in Molecular Biology; Springer US: New York, NY, 2021; pp. 131–138 ISBN 978-1-07-161044-2.

10. Wu, W.; Shang, J.; Feng, Y.; Thompson, C.M.; Horwitz, S.; Thompson, J.R.; Macintyre, E.D.; Thornberry, N.A.; Chapman, K.; Zhou, Y.-P.; et al. Identification of Glucose-Dependent Insulin Secretion Targets in Pancreatic β Cells by Combining Defined-Mechanism Compound Library Screening and SiRNA Gene Silencing. J Biomol Screen 2008, 13, 128–134, doi:10.1177/1087057107313763.

11. Aamodt, K.I.; Aramandla, R.; Brown, J.J.; Fiaschi-Taesch, N.; Wang, P.; Stewart, A.F.; Brissova, M.; Powers, A.C. Development of a Reliable Automated Screening System to Identify Small Molecules and Biologics That Promote Human β-Cell Regeneration. American Journal of Physiology-Endocrinology and Metabolism 2016, 311, E859–E868, doi:10.1152/ajpendo.00515.2015.

12. Szczerbinska, I.; Tessitore, A.; Hansson, L.K.; Agrawal, A.; Ragel Lopez, A.; Helenius, M.; Malinowski, A.R.; Gilboa, B.; Ruby, M.A.; Gupta, R.; et al. Large-Scale Functional Genomics Screen to Identify Modulators of Human β-Cell Insulin Secretion. Biomedicines 2022, 10, 103, doi:10.3390/biomedicines10010103.

13. Bevacqua, R.J.; Dai, X.; Lam, J.Y.; Gu, X.; Friedlander, M.S.H.; Tellez, K.; Miguel-Escalada, I.; Bonàs-Guarch, S.; Atla, G.; Zhao, W.; et al. CRISPR-Based Genome Editing in Primary Human Pancreatic Islet Cells. Nat Commun 2021, 12, 2397, doi:10.1038/s41467-021-22651-w.

14. Grotz, A.K.; Abaitua, F.; Navarro-Guerrero, E.; Hastoy, B.; Ebner, D.; Gloyn, A.L. A CRISPR/Cas9 Genome Editing Pipeline in the EndoC-?H1 Cell Line to Study Genes Implicated in Beta Cell Function. Wellcome Open Res 2020, 4, 150, doi:10.12688/wellcomeopenres.15447.2.

